# Modeling predicts mechanisms altered by mutations of the SARS-CoV-2 delta and omicron variants

**DOI:** 10.1101/2022.02.23.481492

**Authors:** Jason Pearson, Timothy Wessler, Alex Chen, Richard C. Boucher, Ronit Freeman, Samuel K. Lai, Raymond Pickles, M. Gregory Forest

## Abstract

We apply our mechanistic, within-host, *pre-immunity*, respiratory tract infection model for unvaccinated, previously uninfected, and immune-compromised individuals. Starting from published cell infection and viral replication data for the SARS-CoV-2 alpha variant, we explore variability in outcomes of viral load and cell infection due to three plausible mechanisms altered by SARS-CoV-2 mutations of delta and omicron. We seek a mechanistic explanation of clinical test results: delta nasal infections express ∼3 orders-of-magnitude higher viral load than alpha, while omicron infections express an additional 1 to 2 orders-of-magnitude rise over delta. Model simulations reveal *shortening of the eclipse phase* (the time between cellular uptake of the virus and onset of infectious viral replication and shedding) *alone can generate 3-5 orders-of-magnitude higher viral load within 2 days post initial infection*. Higher viral replication rates by an infected cell can generate at most one order-of-magnitude rise in viral load, whereas higher cell infectability has minimal impact and lowers the viral load.

## Introduction

In [1], a mechanistic, within-host, respiratory tract model was developed and applied to predict immediate (hours to days) outcomes of infection and viral load from inhaled SARS-CoV-2 exposures in the respiratory tract. The model assumes all immune system protection is absent on these timescales, consistent with a novel viral exposure, and results were presented based on the best-available information for the 2020 alpha variant. Here we use the model to explore three likely mechanistic effects of the numerous SARS-CoV-2 mutations documented for the delta and omicron variants [2, 3, 4, 5, 6]: (1) *cell infectivity* (probability to infect per infectious virion-cell encounter per second); (2) *duration of the eclipse phase* (the length of time between endocytosis and the shedding of the first daughter virion by a newly-infected cell); and (3) *efficiency of virion replication* (number of infectious virions replicated per day). Our aim is to explore whether, and if so which, variations in the best-known parameter values of the alpha variant can account for the dramatic rises in nasal infectious titers of the delta variant (∼ 3 orders of magnitude) [2, 3, 4, 5] and omicron variant (another ∼ 1.5 orders of magnitude [6]).

We focus this study on the nasal passage only (where there is widely reported testing data of the viral loads for alpha, delta, and omicron). The model outputs the viral load *within* the nasal passage, the flux into the nasopharynx, and the number of infected nasal cells, in the immediate 24-48 hours post infection of one nasal epithelial cell at the entrance of the nasal passage. While these simulations are conducted at the entrance of the nasal passage (i.e., furthest from the nasopharynx), Chen et al. 2022 [1] showed the viral load and infected cell outcomes at the entrance are statistically similar to results from a cell initially infected within the middle two quartiles of the nasal passage.

We model the outcomes in viral load and infected cells across feasible ranges in the kinetic parameters for these 3 mechanistic effects, anchored by results for the alpha variant in Chen et al. 2022 [1]. We refer the reader to Chen et al. 2022 [1] for details of: 1) the physiological details of the nasal passage, including length and circumferential dimensions, thickness and clearance velocity of the mucosal layer, thickness of the periciliary liquid (PCL) layer; 2) the percentage of SARS-CoV-2 infectable epithelial cells (predominantly ciliated cells) in the nasal passage; and 3) the computational model for tracking the spread of viral load and infected cells.

## Results and Predictions

First, we present the total viral load at 1 and 2 days post infection from a single infected cell in the upper nasal passage due to variations in (i) *cell infection probability per encounter per second* and (ii) *replication rate (#/day) of infectious virions from a single infected nasal cell*, for a *fixed 12-hour eclipse phase of the alpha variant* (Figure 1).

**Figure 1:**
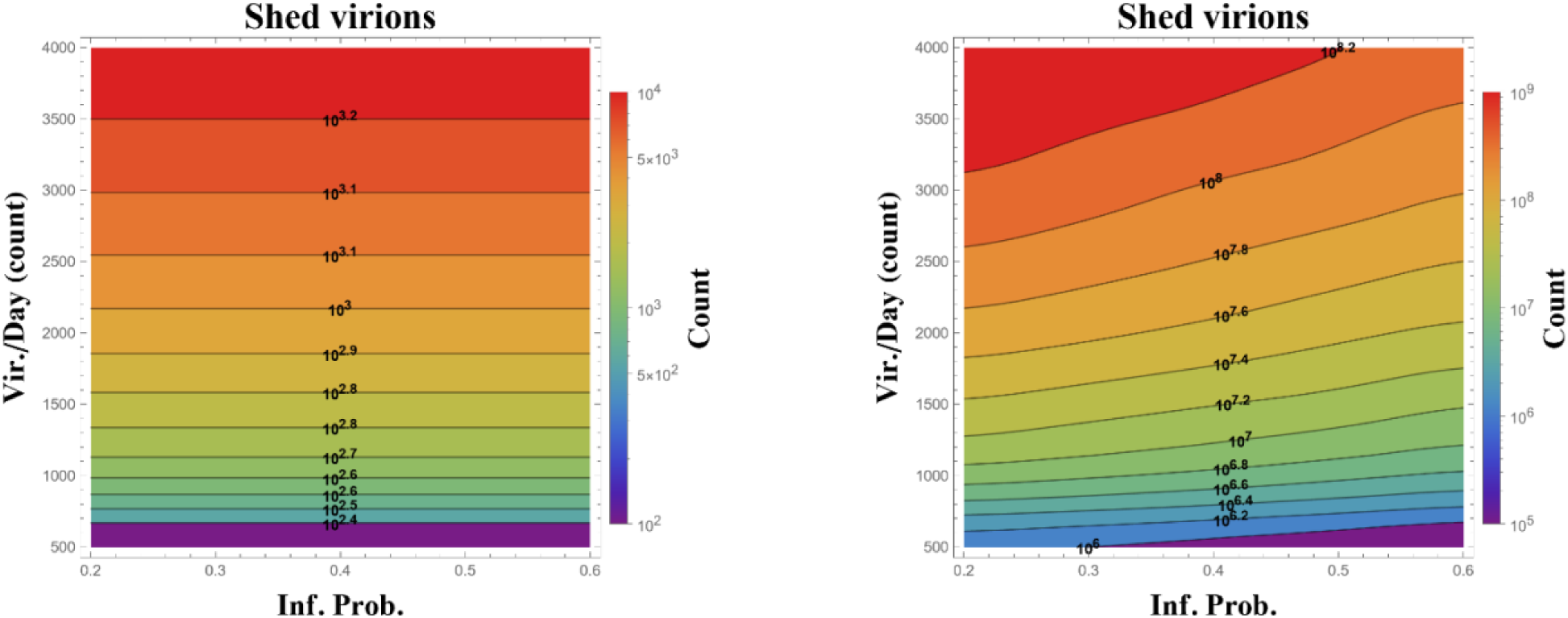
State diagram of total infectious virions replicated over 24 (left) and 48 (right) hours from a single infected (ciliated) cell in the upper nasal passage, calculated over ranges of probability to infect per ciliated cell encounter per second (horizontal) and number of infectious virions replicated per day (vertical). Simulations start at onset of cell infection and assume the alpha variant 12-hour latency time before infected cells begin to replicate. Data presented are averages over 100 realizations, with orders of magnitude on level sets and in the color bar.

### Results from Figure 1

- from [1], the alpha variant, coordinates (0.2, 2000), produces a viral load of ∼ 10^2.9^ after 1 day and ∼ 10^7.7^ after 2 days from one infected cell post infection
- at 1 day the viral load is uncorrelated with probability to infect per encounter-second, then at 2 days is weakly and *negatively* correlated
- these results strongly suggest that spike mutations that *elevate* ACE2 receptor binding alone [2, 3] have minimal impact and if anything lead to even slightly lower viral load after 2 days
- the viral load is positively correlated with the number of infectious virions replicated per day, although the overall contribution toward increased viral load is moderate: after 1, respectively 2, days, viral load rises ∼ 1.0, respectively ∼ 1.5 orders of magnitude by doubling infectious virions per day from 2000 to 4000, and falls ∼ 0.5, respectively ∼ 1.5, orders of magnitude by lowering infectious virions per day from 2000 to 500

*As can be observed clearly in Figure 1, neither one nor a combination of mutations in infection probability and replication rate can account for 3+ orders of magnitude higher viral load relative to the alpha variant*.

### Results from Figure 2 at 24 hours post infection of a single cell in the upper nasal passage

- the shed viral load of the alpha-variant is ∼ 10^3^ for a latency time of 12 hours and 2000 infectious virions shed per day post latency as in Chen et al. 2022 [1]; this shed viral load estimate varies across the state diagram from a minimum of ∼ 10^2.5^ (upper left) to a maximum of ∼ 10^7.5^ (lower right), i.e., 4 orders of magnitude variation
- variations in latency time are dominant: a drop from 12 to 6 hour latency produces a rise of ∼ 3, 3.5, 4 orders of magnitude in shed viral load for fixed 500, 2000, 4000 virions shed per day post latency
- variations in shed virions per day post latency from 500 to 4000 produce ∼ 1.0, 1.5, 2.5 orders of magnitude rise in shed viral load for fixed 12, 9, 6 hour latency
- the infected cell count over the same time course and throughout the state diagram lags the shed viral load by 0.1 (upper left) to 1 (lower right) order of magnitude, and as expected, has strongly correlated parameter gradients with viral load.

Next, we explore the 2-day analog of the 1-day post-infection outcomes in Figure 2, with fixed 0.2 probability to infect per virion-cell encounter, and exploring the potential effects of yet lower latency times. As we incrementally *reduce the latency time from 12 to 2 hours* and run the model over a *2-day period*, there is a dramatic increase in infected cells exiting the eclipse phase and shedding infectious virions. As shown next in Figures 3 and 4, further reduction in latency time dramatically accelerates the viral load over the Figure 2 results. Consequences of the ballistic growth in viral load and infection are long simulation times and large memory requirements. Fast eclipse phases combined with high viral replication require 1 week or longer on the UNC cluster to complete, and require more than 10 TB of memory. In light of these constraints, Figure 3 explores 2 to 12 hour latency times for replication rates of 500-1500 virions per day. Note that one can extrapolate the ∼ 10^7.7^ level set of viral load, indeed all level sets, up to 2000 virions per day, and connect with the results of Figures 1, 2 for the alpha variant.

**Figure 2:**
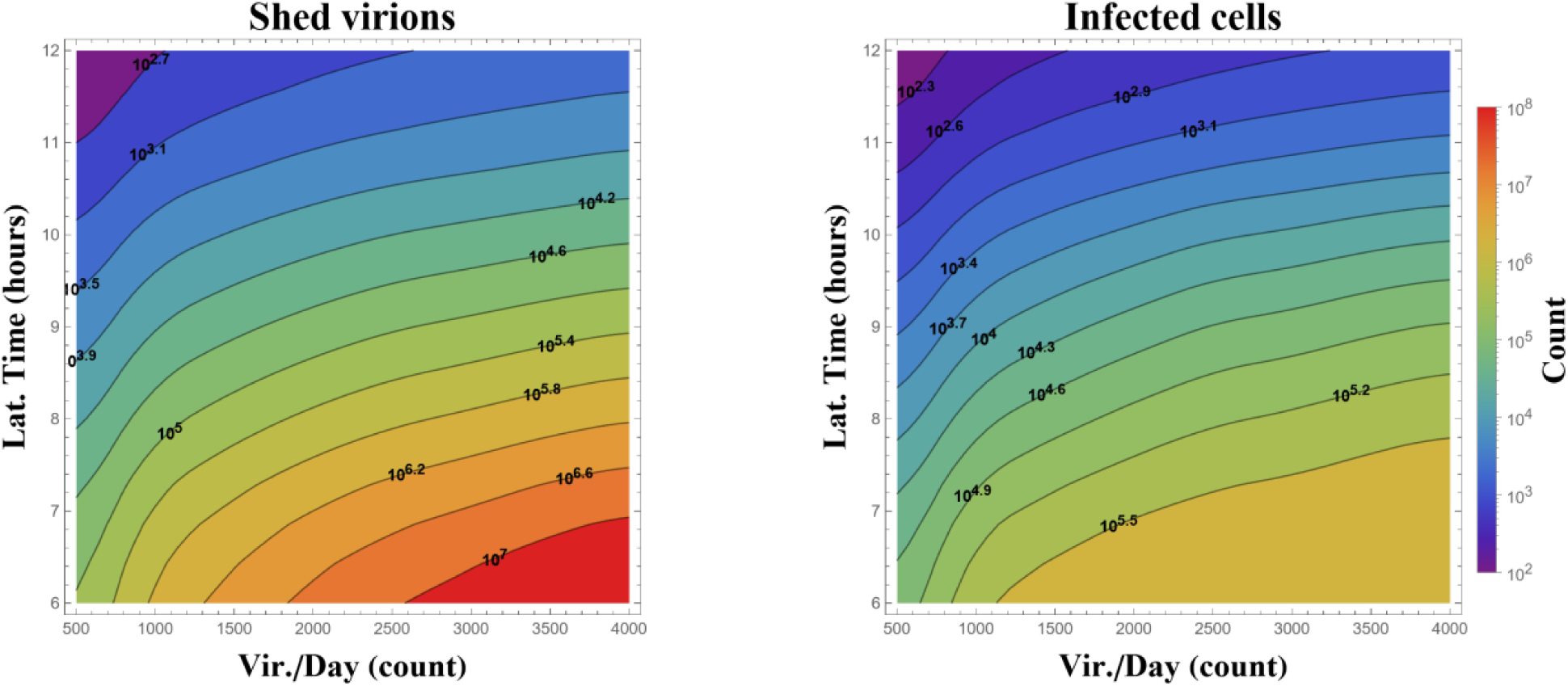
State diagram of *total infectious virions replicated* (left) and *total number of infected cells* (right) at 24 hours post infection from a single infected cell in the upper nasal passage, calculated over ranges of [500,4000] infectious virions shed per day by infected cells (horizontal) and [6,12] hours *latency time* (vertical). Level sets of orders of magnitude are indicated numerically and by color, with all data consisting of averages over 100 simulations, and all simulations are with fixed probability = 0.4 of infection per encounter per sec.

**Figure 3.**
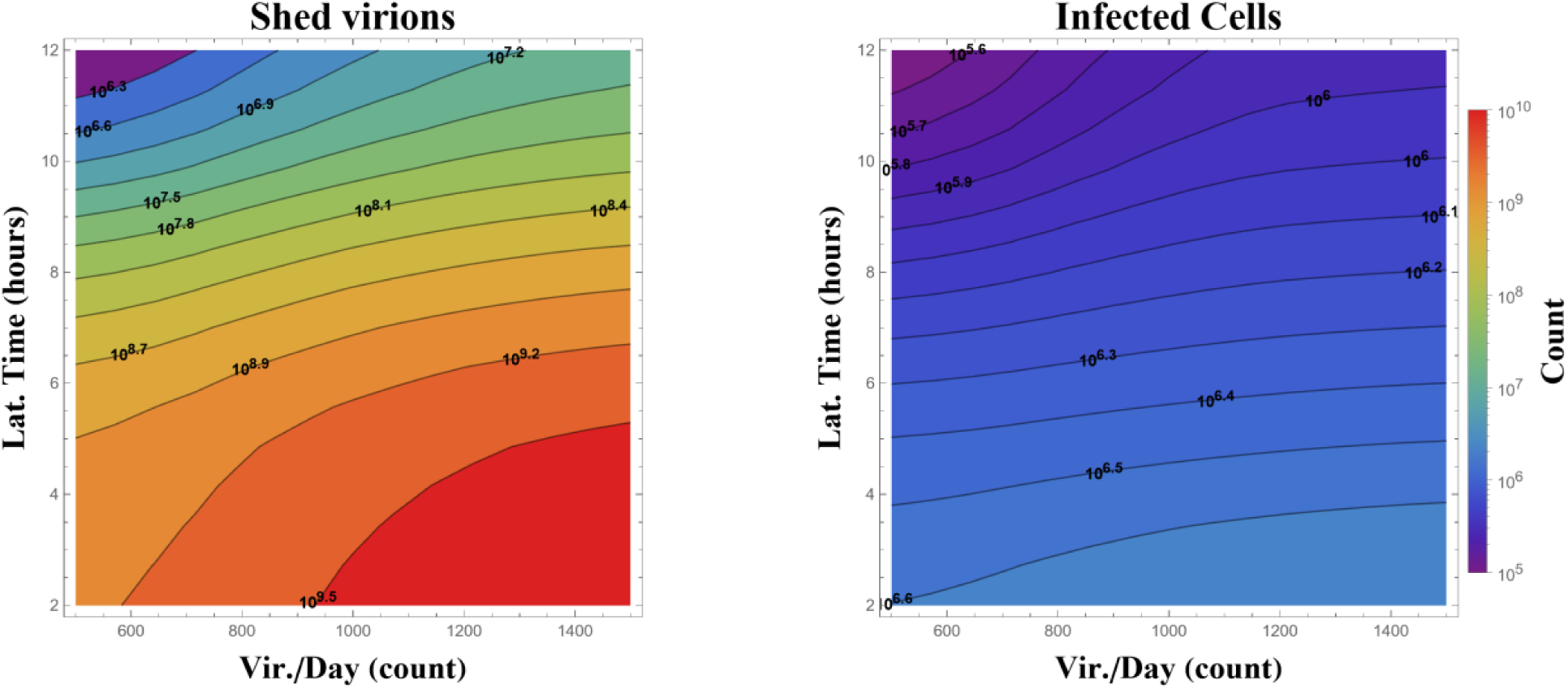
State diagram of total infectious virions replicated (left) and total number of infected cells (right) at 48 hours post infection from a single infected cell in the upper nasal passage, calculated over ranges of [500,1500] *infectious virions shed per day by infected cells* (horizontal) and [2,12] hours *latency time (vertical)*. Level sets of orders of magnitude are indicated numerically and by color, with all data consisting of averages over 100 simulations, and all simulations are with fixed probability = 0.2 of infection per encounter per sec.

**Figure 4.**
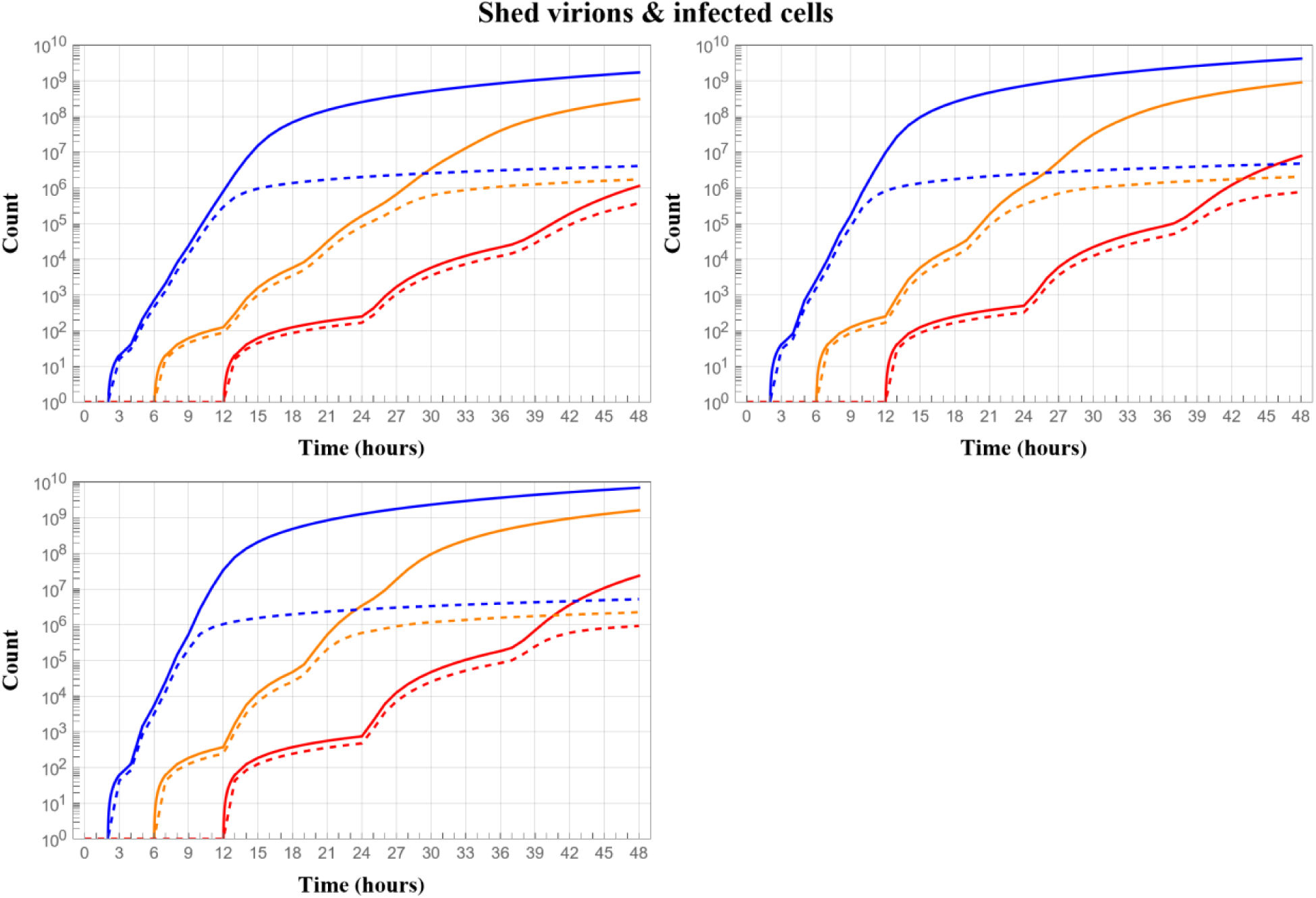
The dynamic evolution of viral load (solid curves) and total infected cells (dashed curves) over two days from a single cell starting from the moment of infection, comparing outcomes for eclipse phases of 12 (red), 6 (orange) and 2 (blue) hours, and for shedding rates of infected cells per day post eclipse phase of 500 (top left), 1000 (top right), 1500 (bottom left). All simulations are for fixed probability 0.2 of infection per cell encounter per sec as used in [Chen et al. 2022] for the alpha variant.

### Results from Figure 3

- the viral load is extremely sensitive to latency time, growing ∼ 2+ orders of magnitude between 12-hour and 6-hour latency, and ∼ 3 orders of magnitude between 12-hour and 2-hour latency, across all fixed values of number of infectious virions per day
- the viral load further grows between 0.5 and 0.9 orders of magnitude by increasing the number of infectious virions per day per infected cell from 500 to 1500;
- the infected cell count grows, but by only ∼1 order of magnitude, lagging significantly behind the total viral load.

These Figure 3 results are consistent with clinical observations of up to 3 orders of magnitude rise in viral titer from the delta variant, albeit still shy of the 4+ orders of magnitude rise in omicron over the alpha variant. By extrapolation of these state diagrams, supported by the robust level sets of viral load, we find *a combination of sufficiently high replication rate (e.g*., *2000 per day) of infectious virions and, more importantly, sufficiently short latency times (e.g*., *2 hours), can reproduce the reported omicron viral loads*.

In Figure 4, we amplify the underlying dynamics over two days of the dramatic impact of reduced latency times for three different shedding rates of infectious virions by infected cells post eclipse phase. We plot single realizations of both the infectious viral load (solid curves) and infected cell count (dashed curves), spanning 48 hours starting from the onset of infection of one cell in the upper nasal passage for latency times of 12, 6, and 2 hours.

### Results from Figure 4

- these single cell realizations illustrate the dynamics underlying the orders of magnitude gains in the state diagrams of Figure 3 by reduced latency times from 12 to 6 to 2 hours
- the curves also show the “episodic surges” in infectious virions, starting from the first passage of latency time as infected cells from the first wave of shed virions emerge from the eclipse phase and begin to replicate, then another surge occurs one latency time later as infected cells from the second wave of shed virions emerge from eclipse phase, and so on
- the shorter the latency time, and as more latency time episodes occur, the episodic surges overlap and smooth out and *appear* to saturate the growth curves, but that appearance is a result of the log plotting scale, as viral load increments are not slowing down.

## Concluding Remarks

The COVID-19 pandemic has been marked by waves of increased population infections and hospitalizations due to emergence of new, apparently more infectious, variants. The delta and omicron variants were responsible for two prominent waves. Reported clinical data for the delta variant revealed ∼3 orders of magnitude increase over alpha in nasal swab viral titers [2, 3, 4, 5], while the omicron variant revealed an additional ∼1.5 orders of magnitude increase [6]. At present, *the mechanisms responsible for such escalations in within-host nasal infection have not been reported*, only documentation of site mutations in spikes and other domains. Three plausible mechanisms of cell infection and viral replication that have become altered by the mutations are explored here, using our recent model of exposure and infection in the respiratory tract.

In Figure 1 we show the mechanistic effect of spike mutations, namely the probability of the virus to bind to cell receptors and thus be taken up by and infect the cell, not only has a very weak effect on the 1-2 day infection, but *stronger binding affinity actually lowers the infectious viral load* generated by an infected cell. Figure 1 further shows the viral load and infected cell consequence of a second plausible mechanistic effect, for the infected cell to be more efficient at replication and shed up to twice as many infectious daughters per day, has at most one order of magnitude impact. These results in Figure 1 assume the 12-hour duration of the eclipse phase for the alpha variant. The upshot of Figure 1 is that higher probability to infect and higher replication rate of infectious daughters by an infected cell cannot come close to reproducing clinical observations of ballistic rises in nasal infections by delta and omicron.

We are thus led to explore consequences of shorter eclipse phases by infected cells. Figures 2, 3, 4 reveal that *lowering the 12-hour eclipse phase of alpha to 6 hours or 2 hours is sufficient to generate 3+ orders-of-magnitude higher viral load in the nasal passage*. We can indeed match 4.5 orders of magnitude rise in viral load of omicron over alpha, but to do so we need to extrapolate from results presented to avoid weeks of simulations and several TB of memory storage on the UNC supercomputing cluster.

